# Potential possibility of using the DNA regions of the internal transcribed spacer ITS and the beta-tubulin gene TUB in the diagnosis of the pathogen *Mycosphaerella linicola* Naumov

**DOI:** 10.1101/2021.01.11.426180

**Authors:** A. Shchukovskaya, O. Skripka, D. Uvarova, A. Shipulin

**Author notes:** Correspondence: A. Shchukovskaya.

## Abstract

During the study, the suitability of the ITS and TUB DNA regions was shown for identifying the target species at the genetic level. However, these DNA regions do not reveal significant differences between the sequences of the target species from different geographic regions. However, when constructing dendrograms of similarities and differences, the sequential differentiation between isolates at the population level was found, which may further affect the results of identification of the target species.

## INTRODUCTION

Pasmo is a dangerous infectious disease of flax caused by fungus *Mycosphaerella linicola*. The pathogen infects cultivated and wild species of the genus flax *(Linum* spp.). The identification of *M. linicola* by classical mycological methods is laborious, time-consuming and not always reliable enough. The advantages of the molecular genetics method are increased reliability, reduced research time, and the use of a small amount of test material. Currently, the most common method for identifying *M. linicola* is a classical polymerase chain reaction followed by sequencing using universal primer systems [Grant L., 2008, Verkley G.J.M. et al, 2013, Verkley G. J. M. et al, 2004]. Of particular interest are the selection of the most effective DNA markers and the assessment of their applicability for identification of *M. linicola.*

## MATERIAL AND METHODS

In this paper the strains of the Pskov and Chinese isolates of the target species were analyzed. Cultural and morphological characteristics of the studied isolates are distinctive of the anamorphic stage of development of M. linicola - Septoria linicola (Spegazzini) Garassini. Further identification of the pathogen was carried out using the molecular genetics method. The identification of the pathogen was based on the determination of the nucleotide sequences of the regions of the transcribed spacer and the beta tubulin gene. Amplification of these loci was carried out with the primers listed in Table 1.

**Table 1.**
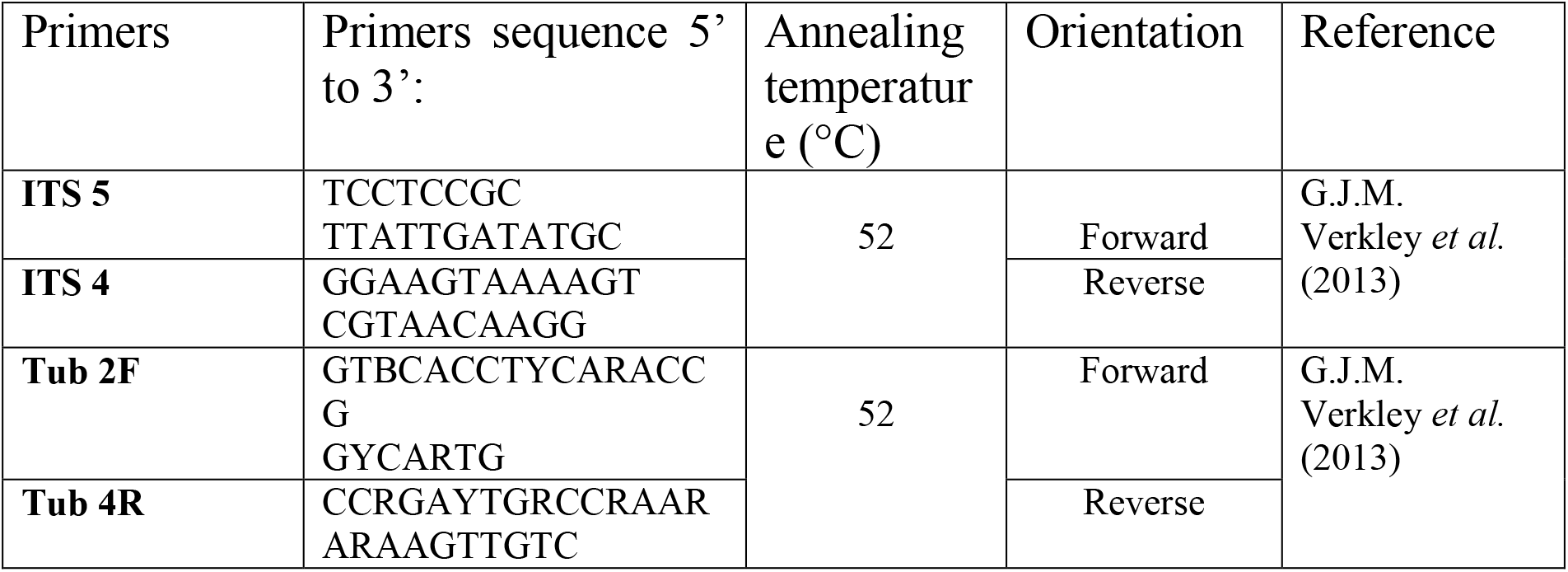
List of primers used in the work

Nucleotide sequences were determined from the obtained PCR products by Sanger Sequencing on a Genetic Analyzer “NANOFOR-05” (ZAO Syntol, Russia). Alignment of nucleotide sequences was performed using the Clustal W algorithm in the BioEdit software.

The original sequences of the target species and nucleotide sequences from the NCBIs GenBank database were used [https://www.ncbi.nlm.nih.gov/genbank]:

✓ Target species (*Septoria linicola*) - Pskov and Chinese isolates (original material) and strain CBS316.37 (NCBI);
✓ *Zymoseptoria tritici* (causing leaf spot of wheat) (family *Mycosphaerellaceae*) strain CBS 398.52, 115943 and IPO320 - to assess variability within the Mycosphaerellaceae family;
✓ *Boeremia exigua* var. *linicola* (causing foot rot of flax) strain CBS 112.28, 113.28 - outgroup.

The neighbor-joining tree was carried out using the MEGA-X program (MEGA-X: Molecular Evolutionary Genetics Analysis version 10.0.5 (Tamura, Stecher, Kumar, 1993-2020).

## RESULTS AND DISCUSSION

Nucleotide sequence analysis of the regions of the internal transcribed spacer (ITS) and the beta-tubulin gene (TUB) revealed a large number of interspecies variable regions, which increases the reliability of recognition of the target species. (Fig. 1, 2)

**Figure 1.**
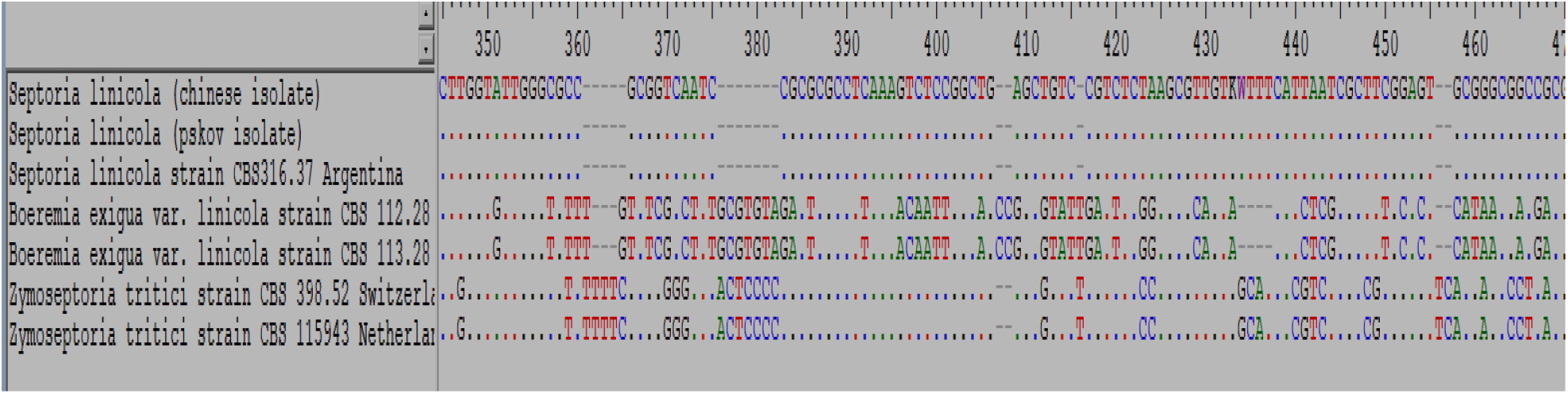
ITS aligned sequences

**Figure 2.**
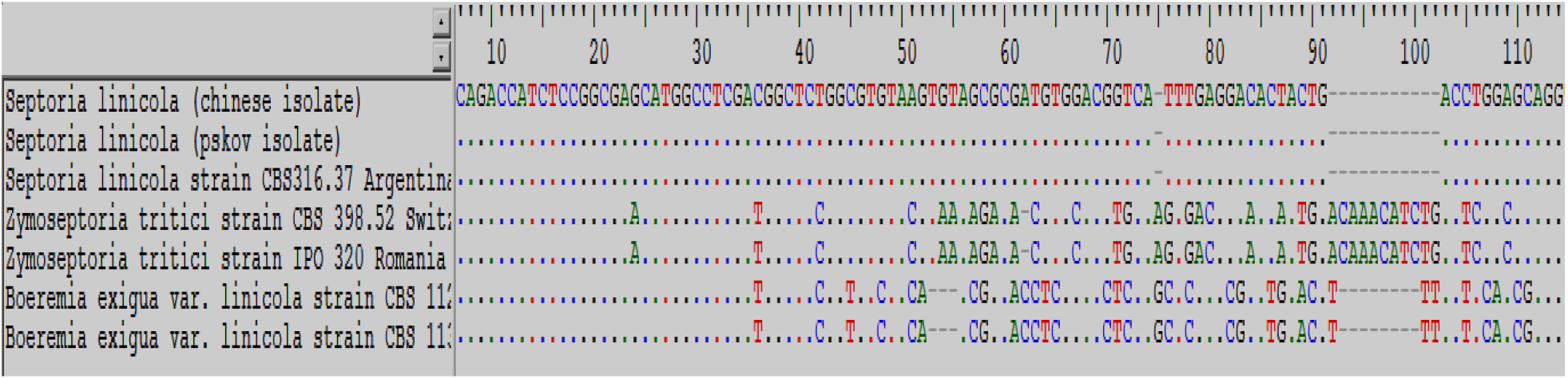
TUB aligned Sequences

However, the analysis of the nucleotide sequences of the target species based on these DNA fragments did not reveal significant differences between them, while according to the results of phylogenetic analysis, *S. linicola* isolates were differentiated at the population level, which did not affect the reliability of the determination of the pathogen of flax (Fig. 1 - 4).

**Figure 3.**
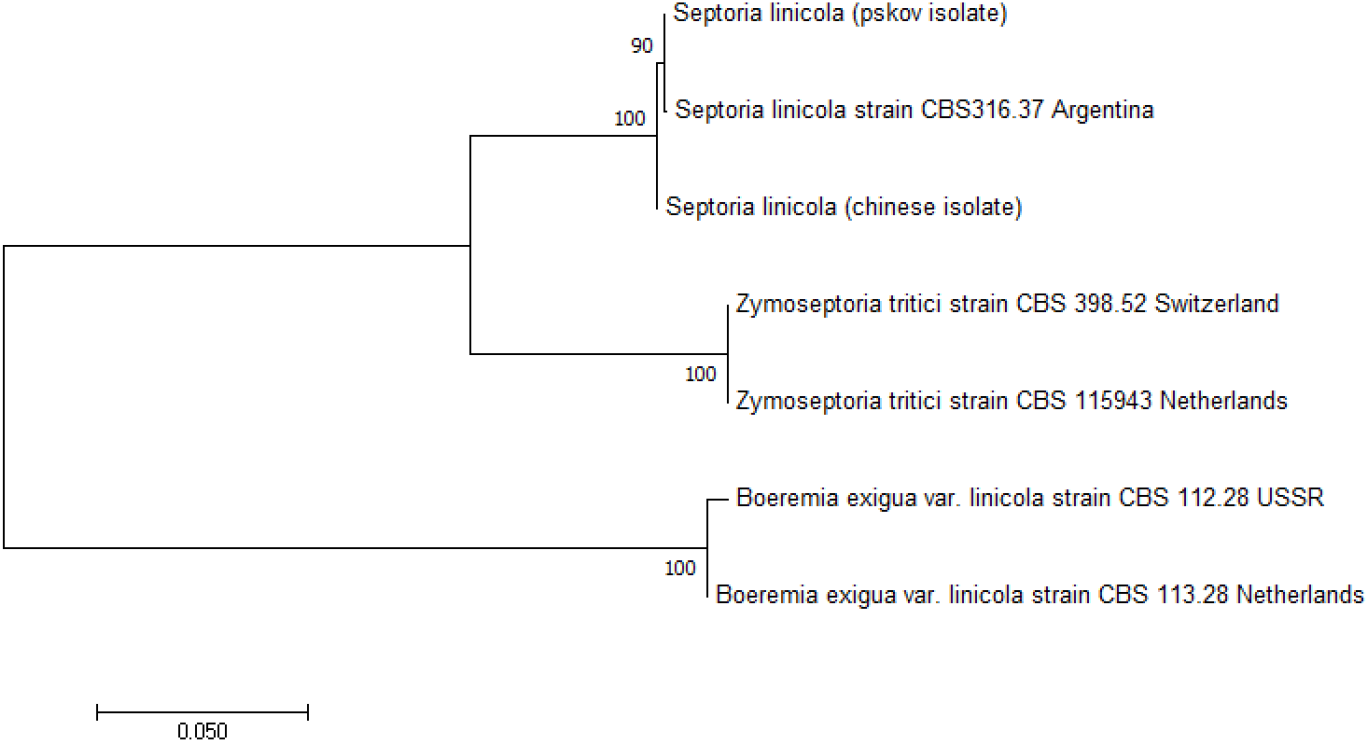
Dendrogram of the sequence similarity of the internal transcribed spacer (ITS) fragment of the target species *Septoria linicola* with the species *Zymoseptoria tritici* and *Boeremia exigua* var. *linicola* (species names and isolate numbers are given), constructed using the nearest neighbor (NJ) linking method. Stability determined by bootstrap analysis (1000 replicates), given as a percentage above the branches

**Figure 4.**
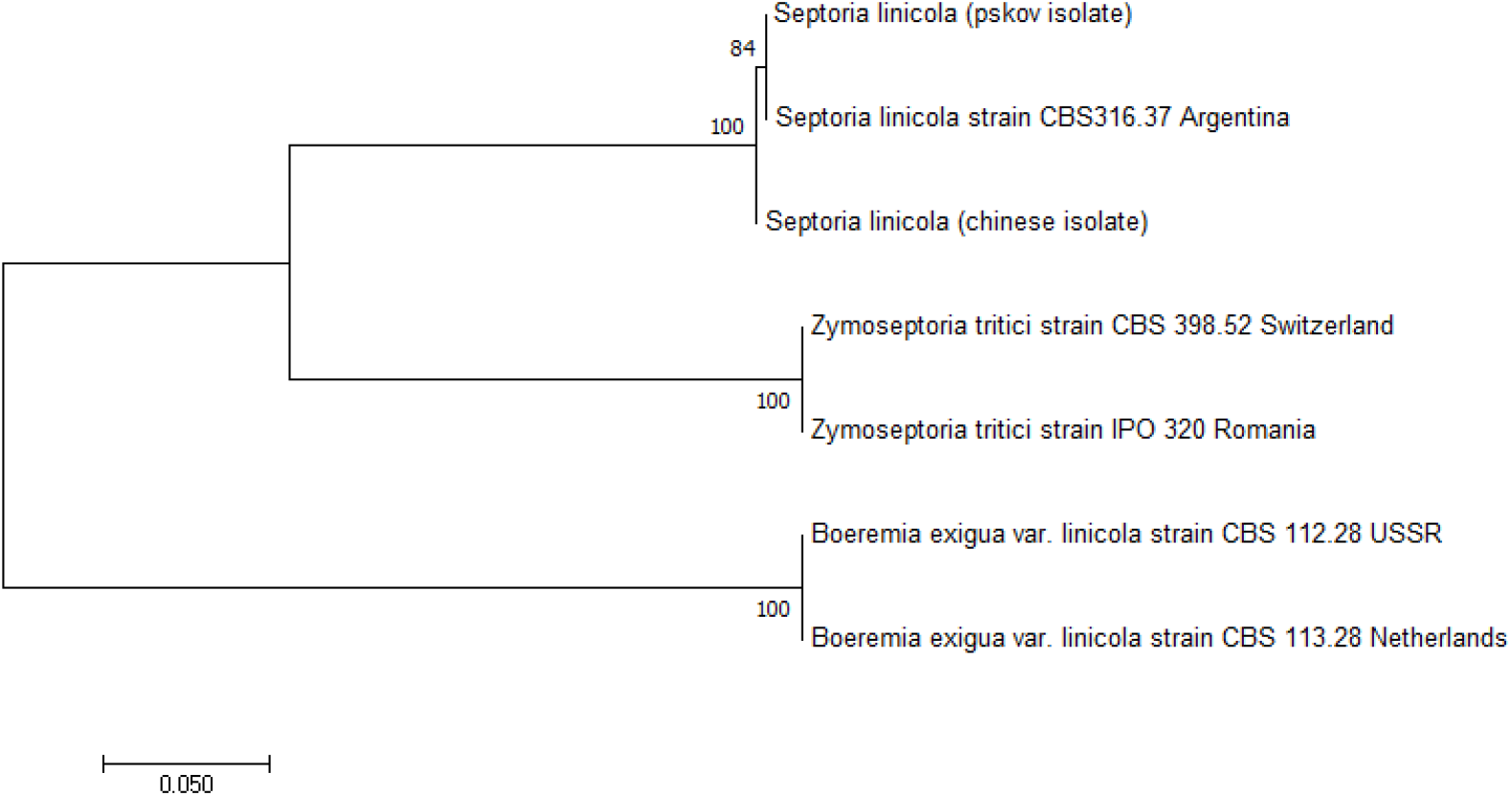
Dendrogram of the sequence similarity of the beta-tubulin (TUB) gene fragment of the target species *Septoria linicola* with the species *Zymoseptoria tritici* and *Boeremia exigua* var. *linicola* (species names and isolate numbers are given), constructed using the nearest neighbor (NJ) linking method. Bootstrap stability (1000 replicates) is given as a percentage over the branches.

Dendrograms of the sequence similarity for fragments of the ITS regions and the beta-tubulin gene TUB show that the value of the bootstrap test for clusters of the target species is 100%, which is sufficient to recognize the existence of these clusters as reliable (Fig. 3, 4). However, within these clusters, differences are observed between the Pskov and Chinese isolates at the population level. The Pskov isolate is similar to the Argentine isolate (the bootstrap test value for the cluster is 90% for a fragment of the ITS region and 84% for a fragment of the beta-tubulin TUB gene region), thus it can be assumed that the Pskov isolate is of Argentine origin and was brought to the territory of the former USSR with seeds.

In its turn, *S. linicola* is well differentiated from *Z. tritici* (family Mycosphaerellaceae) as well as from the pathogen of flax ascochitosis - the outgroup (the value of the bootstrap test for these clusters is 100%, respectively), which allows us to conclude that these DNA areas are effective in distinguishing between the studied species.

Based on the data presented, it can be concluded that the regions of the internal transcribed spacer (ITS) and the beta-tubulin gene (TUB) are characterized by a large number of interspecies variable regions, which in turn makes it possible to determine the target species and separate it from other species at the genetic level.

Dendrograms of similarity for the two studied DNA fragments (ITS, TUB) showed differences between the Pskov and Chinese isolates at the population level. The Pskov isolate is similar to the Argentine isolate, at the same time no significant differences between the *S. linicola* sequences were found, thus it can be assumed that the intraspecies variability of the target species does not affect the result of its identification. However, it is necessary to conduct further studies with cultures of the target species from different geographic regions to study the population variability of the species and assess its impact on the reliability of recognition of the target species.

